# Oxygen supply capacity in animals evolves to meet maximum demand at the current oxygen partial pressure regardless of size or temperature

**DOI:** 10.1101/701417

**Authors:** Brad A. Seibel, Curtis Deutsch

## Abstract

Physiological oxygen supply capacity is associated with athletic performance and cardiovascular health and is thought to cause hypometabolic scaling in diverse species. Environmental oxygen is widely believed to be limiting of metabolic rate and aerobic scope, setting thermal tolerance and body size limits with implications for species diversity and biogeography. Here we derive a quantifiable linkage between maximum and basal metabolic rate and their temperature, size and oxygen dependencies. We show that, regardless of size or temperature, the capacity for oxygen supply precisely matches the maximum evolved demand at the highest persistently available oxygen pressure which, for most species assessed, is the current atmospheric pressure. Any reduction in oxygen partial pressure from current values will result in a decrement in maximum metabolic performance. However, oxygen supply capacity does not constrain thermal tolerance and does not cause hypometabolic scaling. The critical oxygen pressure, typically viewed as an indicator of hypoxia tolerance, instead reflects adaptations for aerobic scope. This simple new relationship redefines many important physiological concepts and alters their ecological interpretation.

One sentence summary: Metabolism is not oxygen limited

## Main

The maximum rate of aerobic metabolism (MMR) is an important measure of physiological performance and fitness that integrates neural, cardiovascular, and metabolic systems. Hill and Lupton(*1*) believed that MMR is limited by oxygen supply, a view still widely accepted today (*2, 3*) (*4*) (*5*). The difference between MMR and the standard or basal metabolic rate (BMR), measured under resting, fasted condition, is thought to represent the absolute aerobic scope (AAS) available for organisms to supply oxygen in support of all aerobic function (*6, 7*). The AAS typically increases with temperature to a peak and then declines at a higher “pejus” temperature. The peak in aerobic scope is believed to represent an optimal temperature while its decline at higher temperatures is thought to result from a failure of oxygen supply.

Similarly, geometric constraints on oxygen uptake and delivery are widely believed to cause hypometric scaling, in which the metabolic rate increases at less than direct proportionality to body mass, and may restrict body size at high temperatures (*8, 9*). Selection for maximum aerobic activity is further believed to be an important driver of the evolution of endothermy in vertebrates(*10*). Oxygen supply capacity is strongly linked to athletic performance, cardiovascular health, temperature and hypoxia tolerance in diverse species including humans with implications for species diversity, abundance, distribution, life history and response to climate change.(*8, 11, 12*), (*13*), (*7, 14*), (*15, 16*)

Despite its ecological and medical importance, oxygen supply capacity is rarely directly measured and the selective pressures acting on it are poorly understood. Here we estimate the physiological oxygen supply capacity (*α*, µmol O_2_/hr/gram/kPa) for 47 species from across the tree of life, including mollusks, arthropods and vertebrates, from marine, freshwater and terrestrial environments (Table S1). We compare *α* derived under two commonly measured, yet distinct, oxygen supply challenges; environmental hypoxia at rest and maximum aerobic exercise. We hypothesize that the physiological oxygen transport system has evolved to supply sufficient oxygen to meet maximum demand at the prevailing environmental oxygen partial pressure (PO_2_). The prevailing PO_2_ is that under which a species’ capacity for activity has evolved, regardless of metabolic rate, body size, or temperature.

To obtain sufficient energy for survival, the O_2_ supplied to an aerobic organism (*S*) must meet or exceed its O_2_ demand as described by the *Metabolic Index*(*15*). At any particular temperature and workload, *S = PO*_*2*_\**α*. Oxygen demand is simply the metabolic rate (MR), here estimated by the rate of oxygen consumption at the same workload and temperature. As ambient oxygen declines, a critical PO_2_ (P_crit_) is reached at which a species’ oxygen supply capacity is fully exploited and below which the basal (resting and fasted) metabolic rate (BMR) can no longer be maintained (Fig. 1). When the environmental PO_2_ reaches P_crit_, *α* can be estimated (*α* = *BMR/P*_*crit*_). In the present dataset, *α* ranges over 2 orders of magnitude, from 0.17 in the cephalopod, *Nautilus pompilius*, to 15.28 in the grasshopper, *Schistocerca americana* (Table S1).

**Figure 1.**
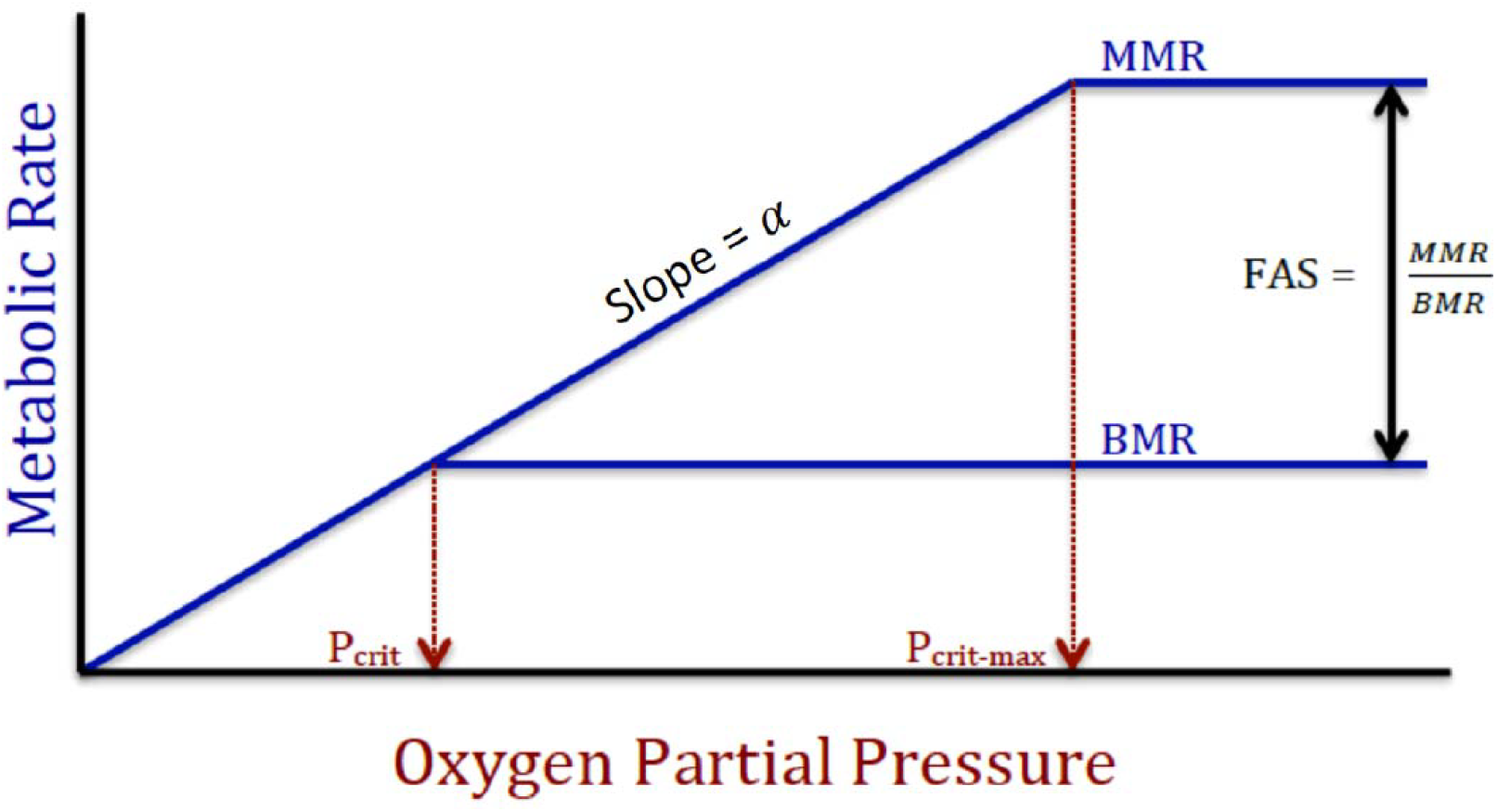
Schematic illustrations of the relationship between metabolic parameters, oxygen and temperature. A) Maximum (MMR) and basal metabolic rates (BMR) and the corresponding critical oxygen partial pressures (P_crit_) are related via the oxygen supply capacity, *α*, which is the slope of the MR-PO_2_ curve.

At MMR, which is typically elicited during intense exercise, *α* is similarly fully utilized and can be independently estimated as *α* =*MMR/P*_*crit-max*_ (the critical oxygen pressure at MMR; Fig. 1). However, P_crit-max_ is rarely measured. We hypothesize that P_crit-max_ is the prevailing PO_2_ in a species’ environment under which their capacity for activity has evolved. To assess the equivalence of *α* under both environmental and physiological oxygen supply challenge, we first estimated *P*_*crit-max*_ = *MMR/α* using the *α* derived from BMR and P_crit_.

Despite more than two orders of magnitude variation in MMR and in *α*, the P_crit-max_ for diverse species is tightly constrained near atmospheric PO_2_. The median value is 19.45 kPa, while the mean is 17.98 ± 5.04 kPa. However, of the 47 species examined, 13 fell below 21 kPa by more than one standard deviation (Fig. 2A). All of those 13 species are known to inhabit persistently hypoxic environments (longer than diel or tidal cycles), including deep-sea oxygen minimum zones, estuaries, poorly ventilated ponds, subterranean burrows and high altitude. While many marine species experience diel or tidal variability in PO_2_ (e.g. intertidal and diel migrating animals), their estimated P_crit-max_ values are within one standard deviation of atmospheric PO_2_ and they are known to employ anaerobic metabolic pathways and metabolic suppression to survive short-term oxygen limitation(*17, 18*). For 9 of the 13 “hypoxic” species, P_crit-max_ has been experimentally determined and it closely matches the predicted values (Table S2). For the additional 4 hypoxic species, the estimated P_crit-max_ values are similar to published environmental PO_2_ values from each species habitat (Table S2). The remaining 34 species (hereafter referred to as “normoxic”) have a mean estimated P_crit-max_ value of 20.62 ± 2.09 and are, thus, expected to achieve maximum activity only at or near atmospheric PO_2_. With few exceptions(*19*), hyperoxia (PO_2_ > 21 kPa) does not result in increases in MMR suggesting that species do not typically evolve the excess supply capacity required to transport the additional oxygen, nor the oxidative capacity to consume any excess oxygen that is delivered to the tissues (Fig. 2B; (*20*)).

**Figure 2.**
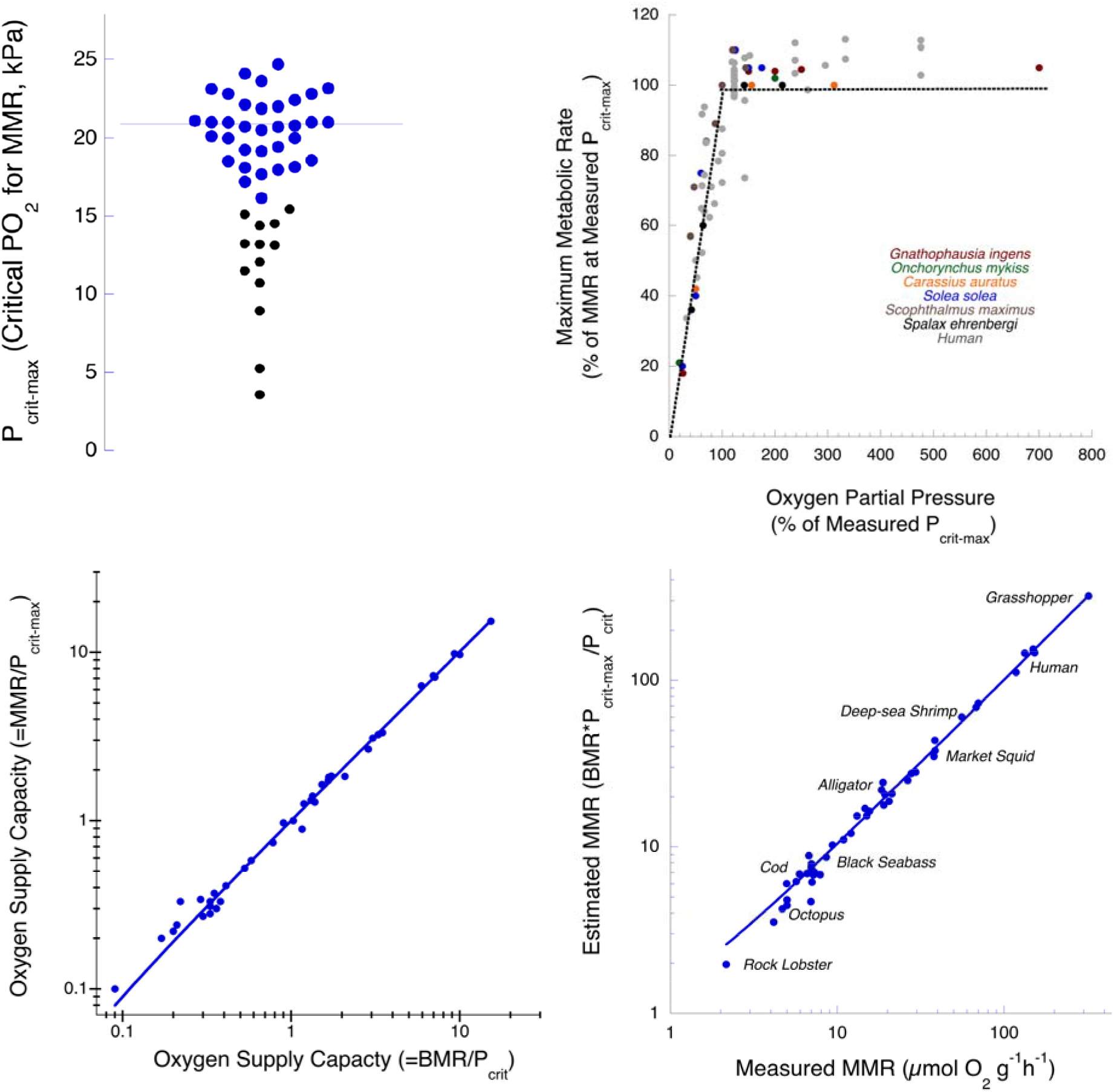
A. Dot plot of P_crit-max_ estimated for the 47 species in this study (Table S1; *P*_*crit-max*_ = *MMR*P*_*crit*_/*BMR*). The mean for all species combined is 17.98 ± 5.04 kPa (n = 47) The black symbols are species for which the estimated P_crit-max_ is below atmospheric PO_2_ (21 kPa) by more than one standard deviation. Those species are all known to inhabit environments with sustained (longer than diel or tidal cycles) hypoxia. Normoxic species (blue symbols) have a mean of 20.62 ± 2.09 (blue line, n = 34). B. The oxygen-dependence of MMR as a percentage of the MMR at P_crit-max_. C. Oxygen supply capacity (*α*) estimated as *BMR/P*_*crit*_ as a function of that predicted as MMR/P_crit-max_ (y = −0.01+ 1X; R_2_ = 1.0; Table S1; Table S2). D. The MMR, predicted as (BMR * PO_2env_)/P_crit_, as a function of the measured MMR (MMR_pred_ = 0.43 + 1MMR_meas_; R^2^ = 0.998; p<0.0001) across diverse species of mollusks, arthropods and chordates.

For the 34 normoxic species, the oxygen supply capacity, estimated as *α* = *BMR/P*_*crit*_, is strongly correlated with that derived as *α* = *MMR/21*, while for hypoxic species, it is correlated with the measured P_crit-max_ or the prevailing environmental PO_2_ (Table S1; Fig. 2C; y = −0.03 + 1.01x; R^2^ = 0.998; p < 0.0001). Thus *α* is equivalent at the limiting oxygen level for any metabolic level (e.g. MMR, BMR or any routine MR between), allowing us to derive a simple relationship between MMR and BMR and their respective critical oxygen pressures (equation 1).

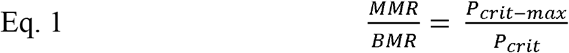

The maximum metabolic rate can be estimated as MMR = *BMR*P*_*crit-max*_/P_crit._ Assuming that P_crit-max_ is air-saturation for normoxic species, and using the estimated environmental PO_2_ and measured P_crit-max_ values for hyoxic species (Table S2), the predicted MMR is precisely correlated with the measured MMR (Fig. 2D; y = 0.26 + 1x; R^2^ = 0.998; p < 0.0001).

The decrement in MMR due to environmental hypoxia can be directly calculated and is proportional to 1/P_crit-max,_ or 4.7% kPa^−1^ for normoxic species. For hypoxic species, P_crit-max_ is lower, which drives a larger relative change in MMR for an equivalent absolute change in PO_2_ (up to 36% kPa^−1^ in *Gnathophausia ingens* living in the oxygen minimum zone in the California Current). Because of the sigmoidal shape of the oxyhemoglobin dissociation curve, the linear relationship between MO_2_ and PO_2_ revealed by our analysis was unexpected. However, we hypothesize that the combined effect of reduced arterial PO_2_ and a right-shift of the oxyhemoglobin curve due to reduced arterial pH during exercise results in a linear decline in MMR with decreasing PO_2_ (Fig. 2C). This is supported experimental work in humans, which demonstrates that arterial PO_2_ decreases linearly with inspired PO_2_ under maximum exercise (*21*). The maximum metabolic rates measured under hypoxia for several fish species(*22–25*) and for the subterranean mole rat(*26*) are all much lower than measurements made at or above P_crit-max_ and are within 10-20% of the values predicted as *α*PO*_*2*_ (Fig. 2B; Table S1). In humans, the measured decrement in MMR with increasing altitude (decreasing PO_2_; (*11, 27*)) also closely matches the predicted reduction (Fig. 2B(*21*)). Human populations adapted to high altitude have a similar MMR but lower P_crit-max_ and higher *α* compared to those at sea level(*28*).

Across both normoxic and hypoxic species, *α* is correlated with both basal and maximum oxygen demand, but the correlation is much stronger with MMR (Fig. 3A). Furthermore, hypoxic species have a higher *α* than normoxic species at a given metabolic rate (ANCOVA; p < 0.001; Fig. 3A). Thus, natural selection acts on the oxygen supply pathways primarily in support of maximum metabolic demand at the prevailing environmental PO_2_, which is P_crit-max_. Athletic performance is achieved by concomitant adjustments in both oxygen demand (i.e. muscle oxidative capacity) and oxygen supply at a given environmental PO_2_.

**Figure 3.**
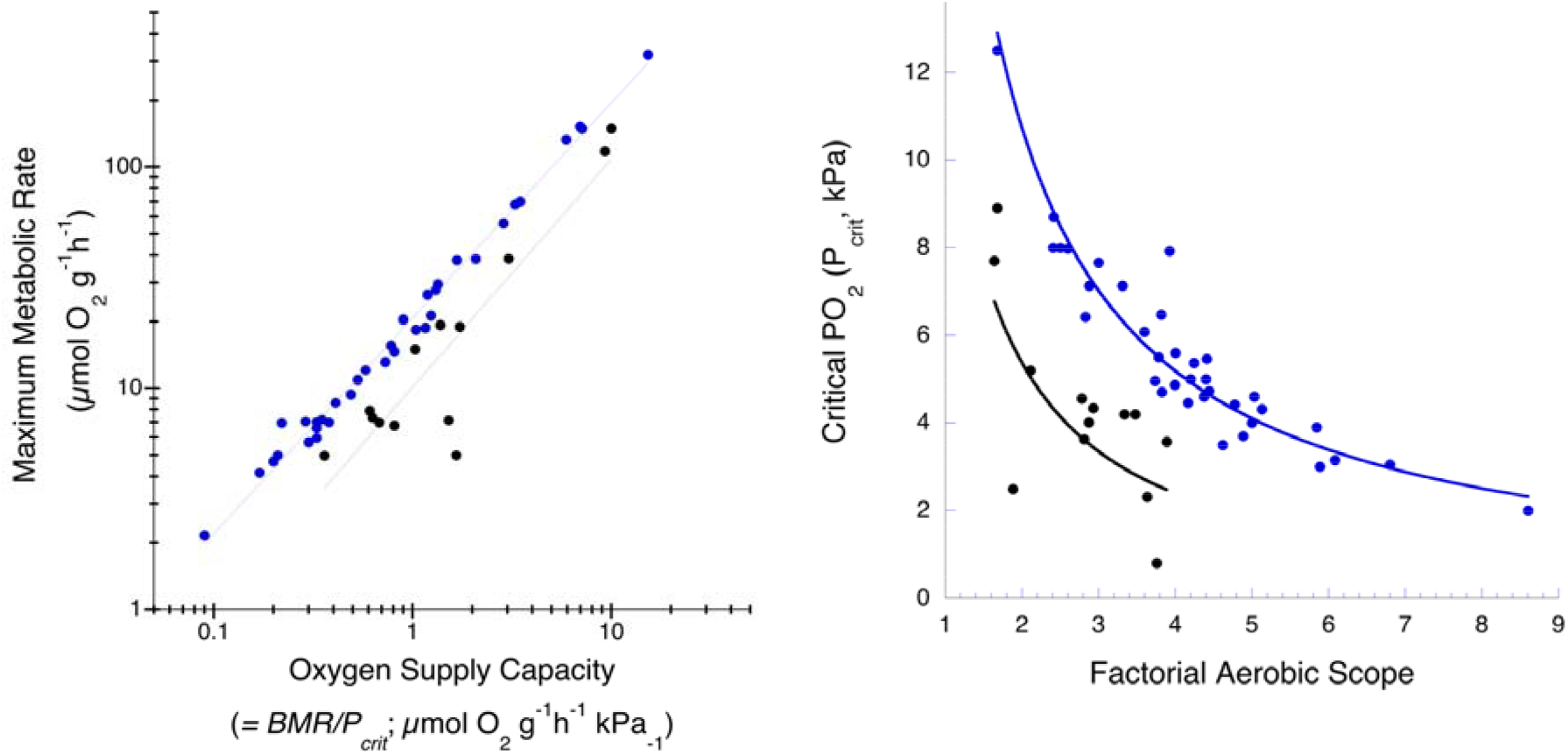
Maximum metabolic rate and critical oxygen levels are related to the oxygen supply capacity (*α*) in normoxic (blue symbols) and hypoxic (black symbols) species. A. MMR as a function of *α* (defined as *BMR/P*_*crit*_; *MMR* = 21 *α*^0.98^; R^2^ = 0.998). For hypoxic species, MMR = 13 *α*^1.0^ (R^2^ = 0.99). B. Factorial Aerobic Scope (FAS = MMR/BMR) as a function of critical PO_2_ (P_crit_ kPa). For normoxic species, FAS = 18 P_crit_^−0.89^ (R^2^ = 0.86). For hypoxic species, FAS = 13P_crit_^−1.0^ (R^2^ = 0.92).

While cardiac output is thought by many to limit MMR, limitation of aerobic performance in hypoxia is typically attributed to gas exchange surface area (lungs or gills) or the oxygen affinity of respiratory proteins(*17, 18, 29*). Despite the many differences between the physiological responses to changing PO_2_ and workload(*30*), the present findings argue strongly that environmental and physiological oxygen supply challenges are met by an equivalent oxygen supply capacity. We suggest that the P_crit-max_ and, thus hypoxia tolerance (see below), is set by the efficacy of oxygen exchange between the environment and the blood (respiratory protein oxygen affinity and gas exchange surface area), while the metabolic rate is primarily determined by the oxygen carrying capacity of the blood (respiratory protein concentration) and cardiac output.

The fact that the oxygen supply capacity is equivalent for species across a range of body sizes and temperatures further argues that *α* evolves and acclimates to meet changing demands with body size and temperature. As a result, not only are MMR, BMR, and their respective critical oxygen partial pressures linked (Eq. 1), but their temperature and body mass scaling coefficients are also mechanistically and quantifiably linked. Temperature (T) and body mass (M) effects on physiological processes or rates can be estimated as *aM*^*b*^ *exp(-E /k*_*B*_ *T)*(*9*), where *a* is the species-specific metabolic rate or P_crit_, *b* is the mass-dependence, *E* is the temperature dependence and *k*_*B*_ is the Boltzmann constant. Inserting these coefficients into eq. 1, we see that *E*_BMR_ = *E*_MMR_ + *E*_Pcrit_ - *E*_Pcrit-max_ and, similarly, *b*_BMR_ = *b*_MMR_ + *b*_Pcrit_ - *b*_Pcrit-max_.

The PO_2_ available in air or in air-saturated water is consistently near 21 kPa regardless of temperature. Thus, P_crit-max_ for normoxic species must be constant across a body size and temperature range (i.e. the temperature and scaling coefficients, *E* and *b*, for P_crit-max_ are zero). Some species may migrate, ontogenetically or on a diel or seasonal basis, across an oxygen and/or temperature gradient. In that case, the evolved P_crit-max_ may change with size or temperature (see below). However, if P_crit-max_ is constant, MMR and *α* have equivalent temperature and mass coefficients at any temperature within the natural range. This suggests that the physiological oxygen supply capacity matches maximum demand at the prevailing PO_2_ regardless of temperature (within the natural range) or body size. Despite nearly a century of research into the effects of both body mass and temperature on metabolism, the data required to test the current relationships is sparse. Furthermore, factorial aerobic scope (MMR/BMR) and P_crit_ must scale with opposite slopes in response to both body mass and temperature (e.g. *b*_MMR_ – *b*_BMR_ = -*b*_Pcrit_).

### Temperature Effects

Only for a handful of species (Table S3) have MMR, BMR and P_crit_ all been measured at more than one temperature. For those species, the temperature coefficients for BMR (*E*_BMR_), predicted as *E*_*MMR*_ + *E*_*Pcrit*_ + E_Pcrit-max_, are strongly correlated with the measured coefficients (Fig. 4A) and the measured MMR correlates with the predicted MMR across the temperature range for each species (Fig. 4B). This suggests that oxygen supply capacity increases with temperature to match increasing metabolic oxygen demand across a temperature range.

**Figure 4.**
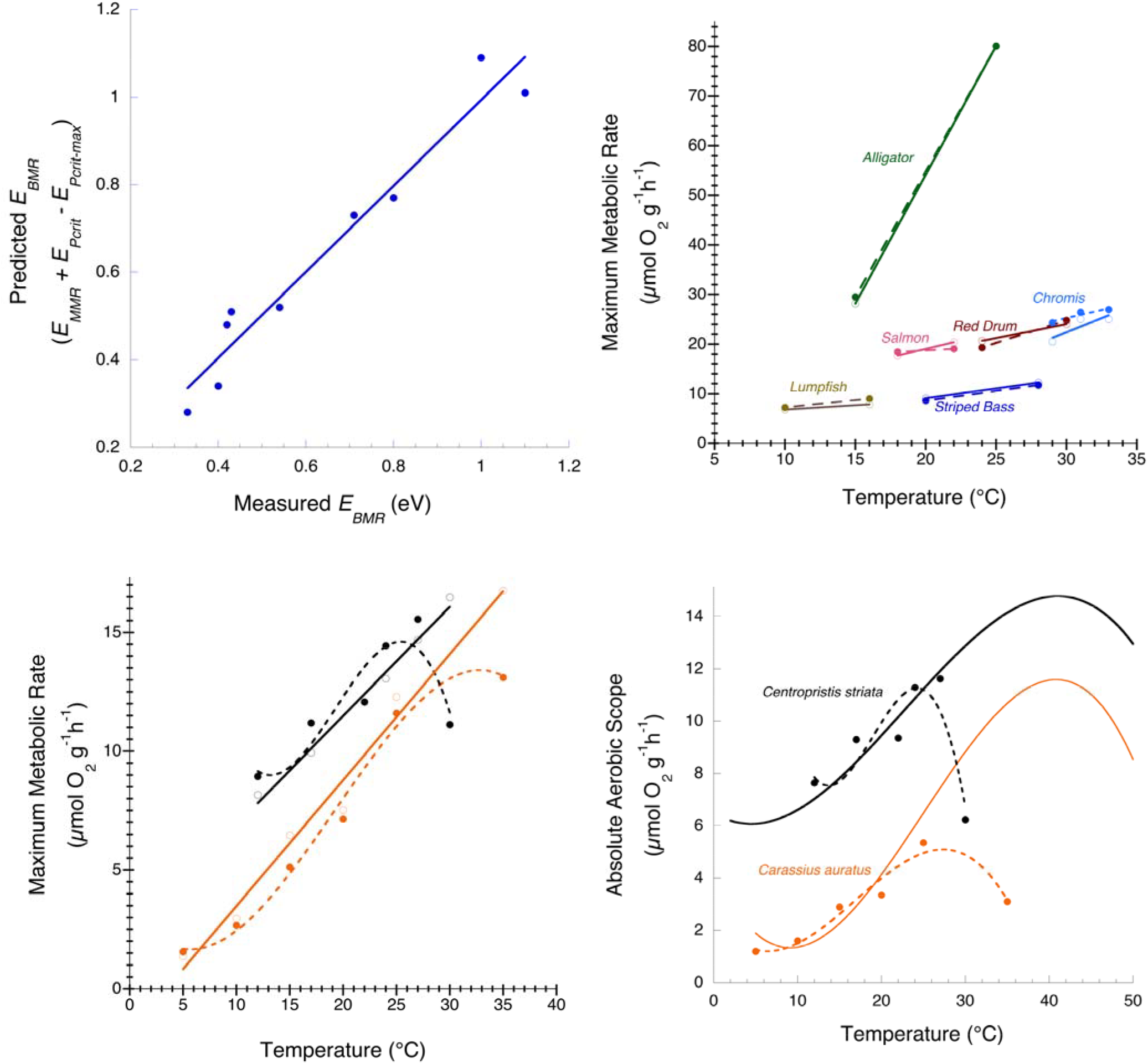
Temperature dependence of measured and predicted MMR and absolute aerobic scope (MMR – BMR) in species for which MMR, BMR and P_crit_ have all been measured at more than one temperature. A) The temperature coefficient for BMR, estimated as *E*_BMR_ = E_MMR_ + E_Pcrit_, as a function of that directly measured (y = 0.01 + 0.98x; R^2^ = 0.94). B) Measured MMR values (solid symbols, dashed lines) and values estimated as *BMR*P*_*crit-max*_/*P*_*crit*_ (open symbols, solid lines). C, D. *Centropristis striata* (Black Seabass) and *Carassius auratus* (goldfish). The measured MMR (C) and aerobic scope (D) closely match predicted values until high temperatures. The decline in MMR at high temperature is not due to a failure of oxygen supply, which would similarly impact P_crit_ and the predicted values. See supplementary tables for data and references.

Aerobic scope (AS), defined as the absolute (AAS = MMR-BMR) or factorial (FAS = MMR/BMR) difference between maximum and resting metabolic rates, is believed to represent the capacity to perform all aerobic activities beyond maintenance metabolism. The pioneering studies of Fry and Hart(*6*), as well as many studies since, show that, MMR and AAS increase with temperature to a peak and then plateau or decline in many species (Fig. 1C; 4C). The temperature at which AAS peaks is widely held as an optimum temperature (T_opt_), while loss of aerobic scope at higher temperatures is often interpreted as an oxygen supply limitation(*7*). Oxygen supply limitation is believed by many (though not all(*31, 32*)) to be an important determinant of temperature and biogeographic limits for ectothermic animals. As such, it has played a prominent role in efforts to predict species’ responses to climate change(*5*).

This “oxygen- and capacity-limited thermal tolerance hypothesis”(*7*) is testable using the new relationship (eq. 1) revealed here. We find that, at temperatures that cause a decrement in the measured MMR, the MMR (and AAS) predicted from BMR and P_crit_ continues to increase in the species for which data are available (Fig. 4C). In other words, there is no decrement in the oxygen supply capacity, *α*, within the measured temperature range. This suggests that the limitation on MMR at high temperature results from something other than oxygen supply. A failure of oxygen supply capacity would affect both P_crit_ and MMR resulting in a decline in both measured and predicted MMR and AAS (Fig. 4C, D).

Absolute aerobic scope is driven almost entirely by changes in MMR(*33*) and, like MMR, it increases with temperature up to a critical temperature and declines at higher temperatures (Figure 1B, C). As such, the increase in AAS with temperature represents increasing costs associated primarily with increasing demands for aerobic activity. The critical thermal maximum (CTmax), defined as the temperature at which BMR equals MMR, P_crit_ equals P_crit-max_, and aerobic scope is nil, is reached only at very high temperatures beyond those experienced by most species. For example, in Black Sea Bass, extrapolation of the temperature curves for BMR, MMR and P_crit_ suggests a peak in AAS above 45°C and a CT_max_ near 65°C (Fig. 5). Thus, the measured peak temperature for AAS (near 24-27°C for Black Sea Bass) is not optimum, but rather is a peak in the metabolic cost to the organism within their evolved temperature range, usually at or just below a critical temperature that elicits complete physiological failure (27-30°C for Black Sea Bass).

**Figure 5.**
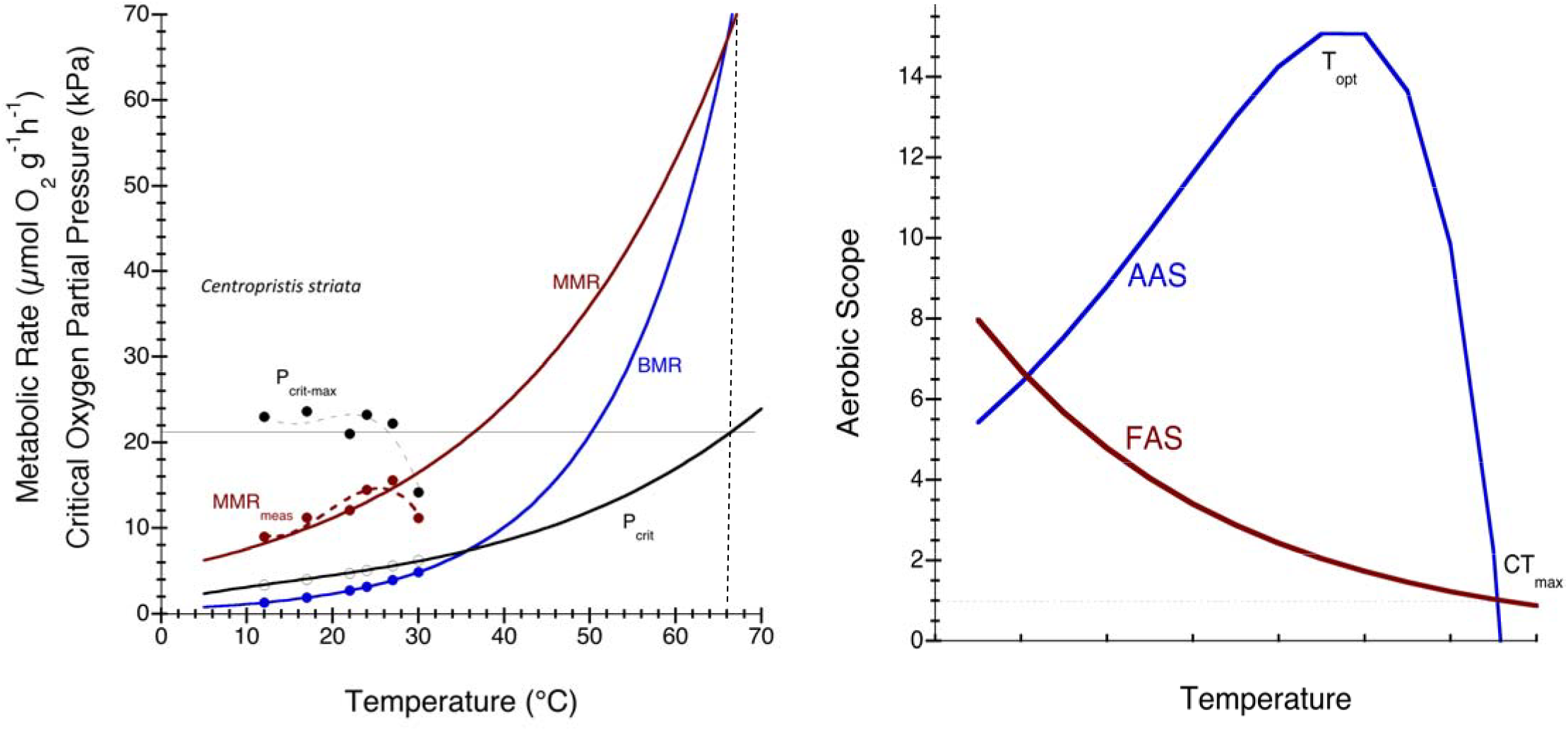
A. Maximum (MMR) and basal metabolic rates (BMR) and the corresponding critical oxygen partial pressures (P_crit_) for Black Sea Bass are shown across temperature range from 12-30°C. The measured temperature coefficients are used to extrapolate to higher temperatures showing that MMR meets BMR, and P_crit_ meets P_crit-max_ (air-saturation), at the traditionally defined critical thermal maximum (CTmax ~65°C), which is well beyond the tolerated temperature range of the species (Slesinger et al., 2019). Note that the measured MMR declines beyond 27°C despite a functional oxygen supply capacity. B. Factorial (*FAS=MMR/BMR*) and Absolute (*AAS=MMR-BMR*) aerobic scope are depicted across a temperature range using the temperature coefficients from Fig. 5A. The CT_max_ is reached where AAS declines to zero, FAS declines to 1, and the P_crit_ equals P_crit-max_. The temperature resulting in a peak in aerobic scope is often considered optimal (T_opt_).

FAS and the Metabolic Index(*15*) are quantitatively equivalent within each species native habitat and, in contrast to AAS, both decline with increasing temperature toward minimal values that are required to support populations. FAS is similar (~3-6) across species due to evolved adjustments in the oxygen supply capacity and in maintenance costs. These adjustments effectively reduce the temperature sensitivity of BMR across species relative to within species. The mean intraspecific E_BMR_ is 0.63 eV while that across species follows E_MMR_ (0.3 eV). The temperature coefficient for FAS is opposite in sign but equal to that for P_crit_ (e.g. E_MMR_ – E_BMR_ = -E_Pcrit_).

### Body mass effects

For most species, P_crit-max_ is constant through ontogeny and will be invariant with body size. For such species, the oxygen supply capacity, *α*, and MMR must scale with identical exponents (eq. 1; *b*_*α*_ = *b*_*MMR*_ + *b*_*Pcrit-max*_). In other words, the oxygen supply capacity increases to meet increasing demand at larger sizes. If P_crit_ is also size-invariant, then BMR must scale with an identical slope to *α*. This undermines a central assumption of the gill-oxygen limitation hypothesis(*8*), which suggests that as ocean warming elevates metabolic rate, fish size will become limited by oxygen supply because the two-dimensional surface area of respiratory organs (i.e. gills or lungs) cannot keep pace with the increasing three-dimensional volume of respiring tissues as fish grow.

The limited data available suggests that P_crit_ is largely size-invariant in fishes(*34*) and insects(*35*). In tropical damselfish, P_crit_ declines slightly with body mass (*b* = −0.1) over several orders of magnitude size range and Pan et al.(*36*) found that P_crit_ *increased* slightly with body mass in the Red Drum, *Sciaenops ocellatus*. Interestingly, Red Drum spend early life in hypoxic estuaries and migrates out to air-saturated coastal waters at larger sizes. Thus, P_crit-max_ likely increases with size for this species. If both P_crit_ and P_crit-max_ are constant, or scale with similar exponents, then *α*, MMR and BMR must scale with similar exponents. FAS scales with a coefficient that is opposite in sign but equal to that for P_crit_ and will thus typically be size invariant (*b*_*FAS*_ = *b*_*MMR*_-*b*_*BMR*_ = -*b*_Pcrit_). Although several studies have found small but significant differences in the scaling coefficients for MMR and BMR, recent work suggests that they scale with similar slopes in fishes and mammals(*37, 38*). Thus, as whole-animal metabolic rate increases with size, the oxygen supply capacity increases to match it. If that were not true (i.e. if *α* did not increase with size in proportion to metabolic demand), P_crit-max_ would increase with size and hyperoxia would be required to exploit the maximum oxidative and muscle capacity. The present findings refute that possibility (Fig. 2B).

### Hypoxia tolerance

A low P_crit_ is usually interpreted as an indication of hypoxia tolerance and, across large PO_2_ gradients, P_crit_ is roughly correlated with habitat PO_2_(*17, 39, 40*). However, P_crit_ must also respond to selection for an increase in aerobic scope, via increased MMR or reduced BMR (eq. 1). In fact, P_crit_ is strongly, inversely correlated with FAS (Fig. 3B). Hypoxic species have a similar range in FAS but a lower P_crit_ at a given FAS (Fig. 3B), suggesting that, rather than hypoxia tolerance *per se*, P_crit_ reflects selective pressure to enhance aerobic scope at the prevailing PO_2_. Without measuring MMR, it cannot be determined whether a low P_crit_ reflects specific adaptation for hypoxia tolerance or elevated aerobic scope. Thus the “incipient limiting oxygen level” (P_crit-max_; (*6, 41*)), which we view as the environmental PO_2_ under which a species is most active and to which the oxygen transport capacity has evolved, is a more useful hypoxia-tolerance metric than P_crit_. P_crit-max_ is the PO_2_ below which a decrement in MMR is certain.

## Discussion

The maximum metabolic rate and aerobic scope are quantifiably linked to resting metabolic rate and critical oxygen pressures. Several recent studies have noted a relationship between metabolic performance and hypoxia tolerance(*15, 20, 22, 25, 42–44*), but the precise equivalency of the oxygen supply cascade during both physical exertion and environmental hypoxia has gone unrecognized. The equivalency revealed here provides the ability to precisely predict the changes in species’ aerobic metabolism with changing PO_2_ and temperature. This simple relationship calls for a conceptual reassessment of aerobic scope, hypoxia tolerance, metabolic scaling and related ecological implications.

For example, in contrast to current thinking(*7*), we suggest that, in many species, the measured decrement in performance at high temperatures results from a failure of the metabolic machinery to use oxygen (e.g. muscle oxidative performance) or an inability of the muscles to produce equivalent work rather than an inability to provide sufficient oxygen. Although, given the evolved match between supply capacity and demand, many physiological systems likely fail at similar temperatures beyond the evolved tolerance range. The exact limitation cannot be known from aerobic scope alone. What becomes clear from the present analysis is that the temperature peak for absolute aerobic scope does not represent an optimum temperature. Similarly, metabolic scaling and temperature-induced reductions in body size cannot result from a size-related oxygen supply limitation as posited by the Metabolic Theory of Ecology(*12*) and the Gill-Oxygen Limitation Theory(*8*) because the oxygen supply capacity evolves to meet increasing demand at large size.

Also in contrast to current thinking, the critical PO_2_ for basal metabolic rate does not specifically reflect hypoxia tolerance. Persistent hypoxia selects for enhanced oxygen supply capacity, which results in a relatively low incipient limiting oxygen level (P_crit-max_). P_crit_ reflects selection for enhanced aerobic scope at a particular environmental PO_2_. Reductions in oceanic PO_2_ have been observed in many subsurface and coastal regions due to warming-induced ocean deoxygenation, upwelling of low oxygen waters and eutrophication. This ocean deoxygenation is projected to accelerate into the future(*45*) (*46*). It is important to note, however, that most shallow marine environments will remain in equilibrium with the atmosphere. While reduced oxygen concentration at constant PO_2_ will not result in reduced metabolic rate (e.g. in most shallow and terrestrial habitats), reduced oxygen partial pressure at depth or due to eutrophication will result in a precise decrement in MMR (1/P_crit-max_; ~5% kPa^−1^ for most shallow-living and terrestrial species), with consequences for vertically mobile and mesopelagic species(*39*).

The ecological and fitness implications of small hypoxia-induced decrements in MMR or aerobic scope (~5% kPa^−1^) are difficult to know, but may impair predator-prey interactions or restrict growth and reproduction(*5*). A recent analysis for marine animals demonstrates that population limits occur below a mean critical value of the Metabolic Index (equivalent to factorial aerobic scope) of ~3(*15*). In other words, a factorial aerobic scope of ~3 is required to support a population and an environmental PO_2_ ~3 times higher than the critical PO_2_ is required to support that aerobic scope. Evolutionary adjustments in oxygen supply capacity and in maintenance costs result in similar FAS (~3-6) in species across the temperature range occupied by animals.

Some reduction in the capacity for aerobic activity will occur for any species exposed to an O_2_ partial pressure below their P_crit-max_. Increasing temperature within a species natural range also elevates metabolic demand, with BMR increasing faster than MMR, leading to reduced FAS. However, as long as the PO_2_ remains at or above P_crit-max_, the evolved maximum metabolic capacity at the higher temperature can be fully realized. Providing excess oxygen (beyond P_crit-max_) will not elevate aerobic scope toward values achieved at lower temperatures. Temperature and hypoxia stress are linked, but their consequences for metabolism and aerobic scope are not equivalent. Acute changes in temperature result in a change to the oxygen supply capacity whereas changes to oxygen partial pressure do not.

In summary, we find that strong selective pressure acts on the oxygen supply system to meet the maximum oxygen demand, despite wide interspecific, temperature- and size-related variation. This finding is consistent with “symmorphosis”, a concept in which each step in the oxygen supply cascade has evolved in concert, without a single rate-limiting step(*47*)(*48, 49*). Proponents of this view suggest that organisms possess little or no excess capacity for oxygen supply nor for its use.

The oxygen supply capacity matches MMR across a size and temperature range (Fig. 3A). The selective pressure on *α* is enhanced for species living in persistent hypoxia, allowing them to achieve metabolic rates similar to those of species living at atmospheric PO_2_, but with a reduced incipient limiting oxygen level(*6*) (P_crit-max_). However, enhanced oxygen supply capacity at a given environmental PO_2_ serves only to elevate MMR. Because BMR and MMR are linked, enhanced aerobic scope must be achieved via efficiency adaptations to reduce maintenance costs thereby reducing BMR relative to MMR. The P_crit_ for BMR is a simple consequence of those balancing selective pressures and is not under direct selection for hypoxia tolerance. Absolute aerobic scope primarily mirrors MMR and does not provide an obvious additional fitness benefit. Species do not evolve excess capacity to supply oxygen nor excess capacity for its use. The traditional views of aerobic scope and hypoxia tolerance are refined in this light. The clear and quantifiable connection between MMR, BMR and P_crit_ provides new power to test and predict the response of animals to changing environmental conditions.

## Author contributions

B.A.S. conceived of and wrote the manuscript. B.A.S and C.D. developed the theory, performed the computations, verified and discussed the analytical results and contributed to the final manuscript.

## Data availability

All data is available in the manuscript and supplementary materials.

## Supplementary Materials

### Methods

The physiological parameters, MMR, BMR and P_crit_, are laboratory measurements taken from published literature. The original studies are given in Tables S1–S3. The measurements range over 5 orders of magnitude in body mass, span nearly the entire range of habitable temperatures, and are from 3 phyla (Arthropoda, Chordata, and Mollusca). The data compiled here are derived from studies using diverse methodologies as appropriate for the diversity of taxa represented. We included all available taxa for which all three physiological metrics have been measured, regardless of the methods employed (e.g. closed, flow-through or intermittent flow respirometry) or the acclimation period allowed. Maximum metabolic rate (MMR) for all species is achieved during or just after exercise protocols induced either via treadmills or swim flumes or following an exhaustive chase. Basal metabolic rate (BMR) is the lowest rate achieved by a particular species at a specified temperature, typically in a resting, fasted state. The critical oxygen partial pressure (P_crit_) is defined as the oxygen partial pressure below which the basal rate of metabolism can no longer be maintained.

Despite the variable methods used, the relationship from equation 1 is extremely robust as indicated by the precision with which MMR is estimated (Fig. 2D). Deviation from the relationship described here may result from 1) a failure to experimentally achieve a truly maximum metabolic rate, 2) a decrement in oxygen demand relative to supply capacity, for example, at temperatures beyond the tolerated range, 3) inaccurate estimate of P_crit-max_, or 4) error in the P_crit_ determination. Several studies note, for example, discrepancies between the MMR achieved by fishes during excess post-exercise oxygen consumption (EPOC; i.e. the “chase” method) and that obtained during maximum sustained swimming using a flume(*52*)(*31*).

Similarly, most of the variation in P_crit_ determination (e.g. between studies of the same species) results from variation in measured BMR, rather than in the determination of P_crit_ itself. Such error most often results from organismal stress or spontaneous activity leading to an elevated rate that is better termed a “routine rate” (RMR). Any elevation in BMR will lead to an underestimated aerobic scope. However, BMR and P_crit_ covary along the MO_2_-PO_2_ curve (the slope of which is *α*, the oxygen supply capacity). Thus, an accurate MMR can be calculated from any paired MR and P_crit_ measurement. Assuming a value for P_crit-max_ that exceeds the true P_crit-max_ (the environmental PO_2_ under which maximum capacity has evolved) will lead to an overestimate of MMR.

The temperature dependence (*E*) of each metabolic metric was determined from the slope of the linear regression of 1/k_B_T versus the natural log of the metric (T in °Kelvin and k_B_ is the Boltzmann constant; Table S3). The temperature and body mass dependencies for each metric are related according to equation 2.

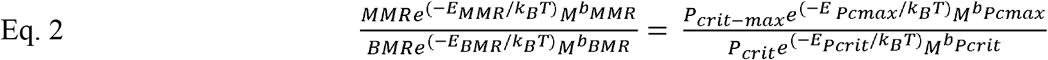

**Table S1:**
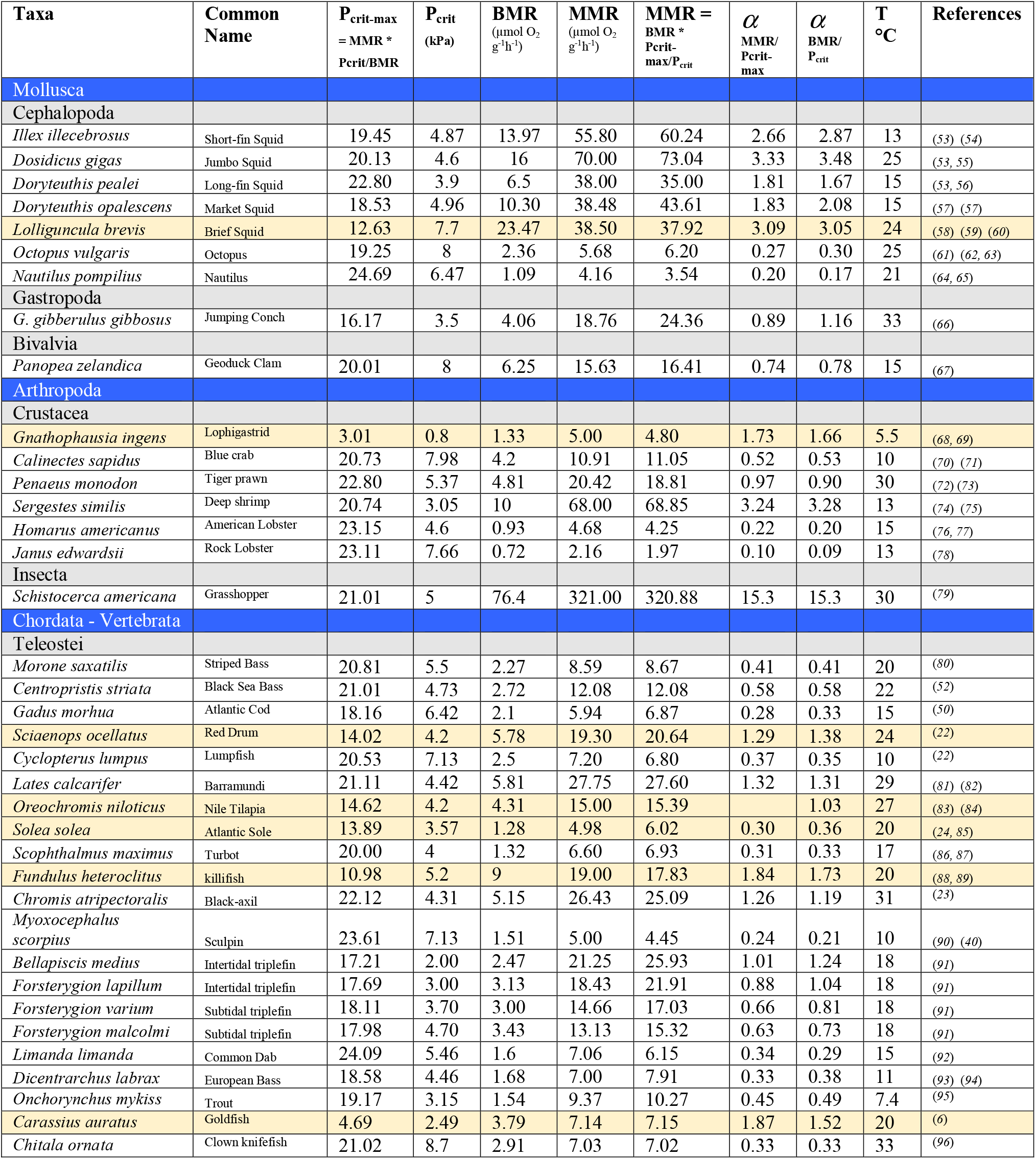
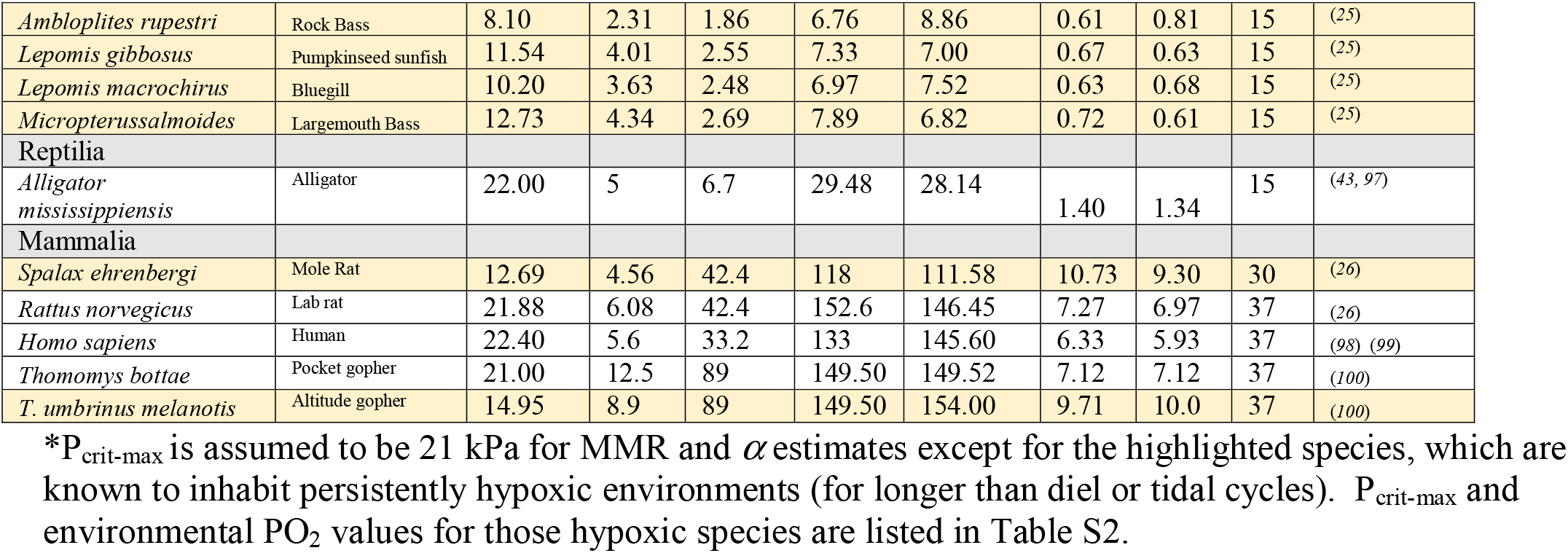
Metabolic measurements for diverse species.

**Table S2.** P_crit-max_ and environmental PO_2_ (PO_2env_) in diverse species.

| Table S2. $P_{crit-max}$ and environmental $PO_2$ ($PO_{2env}$) in diverse species. | | | | |
| --- | --- | --- | --- | --- |
| Species | Predicted<br>$P_{crit-max}$<br>(kPa)<br>$= \frac{MMR * P_{crit}}{BMR}$ | $PO_{2env}$ or<br>Measured<br>$P_{crit-max}$ | Description of $PO_{2env}$ | Ref |
| *Lolliguncula brevis* | 12.63 | 12.44 | Summer $PO_2$ in Chesapeake Bay when squids are present (4 mg/l at 26°C). | (101) |
| *Gnathophausia ingens* | 3.01 | 2.89 | $P_{crit-max}$ is 2.89 kPa (0.95 ml/l at 5.5°C), approximate $PO_2$ maximum in California Current between 400-800 m depth. | (68) |
| *Sciaenops ocellatus* | 14-15.48 | 15 | $PO_2$ high of 15 kPa (5 mg at 20°C) in estuarine habitat. | (102) |
| *Solea solea* | 13.89 | 16.8 | $P_{crit-max}$ is 16.8 kPa. The bottom waters of the Adriatic Sea are known to be hypoxic but a precise figure could not be determined. | (85) |
| *Fundulus heteroclitus* | 10.98 | 10.3 | $PO_2$ range day to night is ~6.7 to 10.3 kPa | (103) |
| *Carassius auratus* | 4.69 | 3.96 | $P_{crit-max}$ (20°C) is 3.96 kPa (Incipient lethal oxygen level). Goldfish ponds are also known to be hypoxic but highly variable. | (6) |
| *Oreochromis niloticus* | 14.62 |  | Highly variable hypoxic environments. | (51) |
| *Ambloplites rupestri* | 8.40 | 11 | $P_{crit-max}$ estimates consistent with MMR measurements at 6 and 9 kPa. | (25) |
| *Lepomis gibbosus* | 11.54 | 11 | $P_{crit-max}$ estimates consistent with MMR measurements at 6 and 9 kPa. | (25) |
| *Lepomis macrochirus* | 10.20 | 11 | $P_{crit-max}$ estimates consistent with MMR measurements at 6 and 9 kPa. | (25) |
| *Micropterus salmoides* | 12.73 | 11 | $P_{crit-max}$ estimates consistent with MMR measurements at 6 and 9 kPa. | (25) |
| *Spalax ehrenbergi* | 12.69 | 11 | 11 kPa is approximate $P_{crit}$ for MMR. Mole rats are known to inhabit hypoxic underground tunnels. | (26) |
| *T. umbrinus melanotis* | 14.95 | 15.4 | Collected at 3200 m = 15.4 kPa. Measured $P_{crit-max}$ is 14.95. | (100) |

**Table S3.**
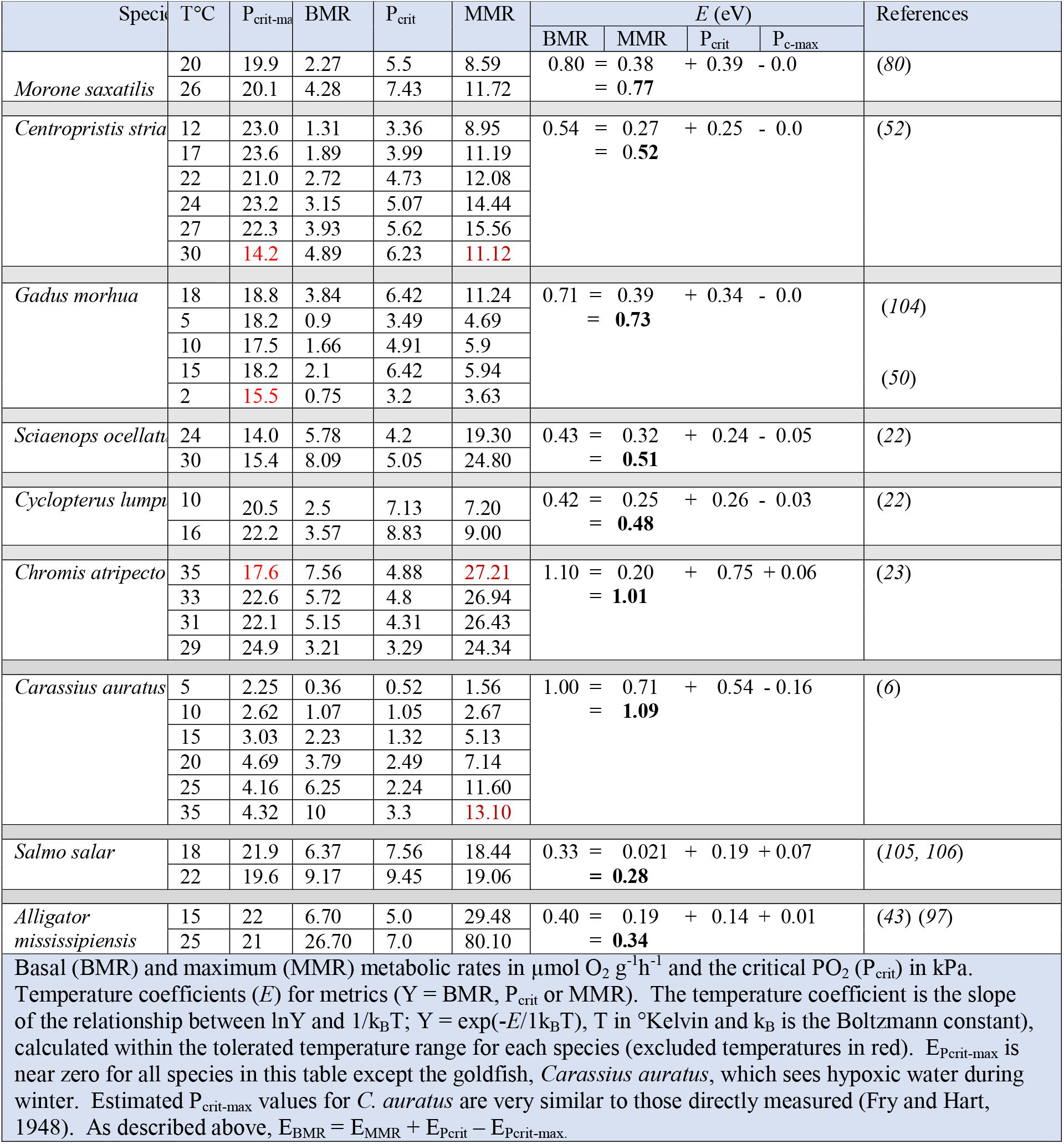
Temperature effects on metabolic rate and critical oxygen pressures.

